# Prediction of Essential Binding Domains for the Endocannabinoid N-Arachidonoylethanolamine (AEA) in the Brain Cannabinoid CB1 receptor

**DOI:** 10.1101/2020.02.19.956003

**Authors:** Joong-Youn Shim

**Affiliations:** Department of Physical Sciences, School of Science, Technology and Mathematics, Dalton State College, Dalton, Georgia, United States of America

## Abstract

∆^9^-tetrahydrocannabinol (∆^9^-THC), the main active ingredient of *Cannabis sativa* (marijuana), interacts with the human brain cannabinoid (CB1) receptor and mimics pharmacological effects of endocannabinoids (eCBs) *N*-arachidonylethanolamide (AEA) and 2-arachidonoylglycerol (2-AG). Given recent intriguing findings that some allosteric modulators can enhance selectively the AEA-activated CB1 receptor, it is imperative to determine the structure of the AEA-bound CB1 receptor. However, due to its highly flexible nature of AEA, establishing its binding mode to the CB1 receptor is elusive. The aim of the present study was to explore many possible binding conformations of AEA within the binding pocket of the CB1 receptor that is confirmed in the recently available X-ray crystal structures of the agonist-bound CB1 receptors and predict essential AEA binding domains. We performed long time molecular dynamics stimulations of plausible AEA docking poses until its receptor binding interactions became optimally established. Our simulation results revealed that AEA favors to bind to the hydrophobic channel of the CB1 receptor, suggesting that the hydrophobic channel holds essential significance in AEA binding to the CB1 receptor. Our results also suggest that the H2/H3 region of the CB1 receptor is an AEA binding subsite privileged possibly over the H7 region. The results of the present study contribute to identifying the (hidden) allosteric site(s) of the CB1 receptor in our immediate future study.

## Introduction

∆^9^-tetrahydrocannabinol (∆^9^-THC), the main active ingredient of *Cannabis sativa* (marijuana), interacts with the brain cannabinoid (CB1) receptor and elicits a wide range of neurological, psychological and biological effects [1]. Continuous marijuana use may increase risks of addiction, chronic pain, mood disorders, and birth defects [2,3].

Recently determined X-ray crystal structures of the CB1 receptor in complex with various ligands [4-7] have revealed the detailed receptor interactions with the bound ligand. Toward understanding molecular mechanisms of marijuana action, these X-ray crystal structures have also shed light on how the ligand activates the receptor upon binding at the molecular level. It is seen in the X-ray crystal structure of the classical cannabinoid full agonist AM11542-bound CB1 receptor [6] that the dimethyl heptyl (DMH) chain of the ligand binds the hydrophobic channel, disrupting the toggle switch of Phe200/Trp356. The toggle switch has been proposed to be required for CB1 receptor activation [8]. The hydrophobic channel appears to be conserved not only in the CB1 receptor but also in other related lipid receptors [9,10]. The classical cannabinoid ∆^9^-THC is expected to bind the CB1 receptor in a way similar to AM11542. Thus, it is likely that the known partial agonistic activity of ∆^9^-THC [11,12] is attributed to its pentyl tail moiety that binds the hydrophobic channel but not as tightly as the DMH chain of AM11542.

Just like Δ^9^-THC, endogenous lipid ligands such as *N*-arachidonylethanolamide (AEA) (Fig 1) and 2-arachidonoylglycerol (2-AG), known as endocannabinoids (eCBs), also interact with the CB1 receptor. AEA was isolated from porcine brain [13] and 2-AG from canine intestines [11]. These eCBs are known to be produced only when biologically demanded [14,15]. A common structural feature of eCBs is a long lipid chain containing the polyene linker moiety and the pentyl tail moiety (Fig 1), which makes them extremely flexible and allows them to adopt millions of conformations. Identification of the bioactive conformation of AEA to the CB1 receptor can be quite elusive due to its potential to adopt many low-energy binding conformations only a few of which would be responsible for receptor activation. AEA is a partial agonist at CB1 receptors [16,17] just like Δ^9^-THC but somewhat more potent than Δ^9^-THC in activating the CB1 receptor [1]. Without any known X-ray crystal structure of the AEA-bound CB1 receptor, the nature of binding interactions of AEA with the CB1 receptor remains poorly understood.

**Fig 1.**
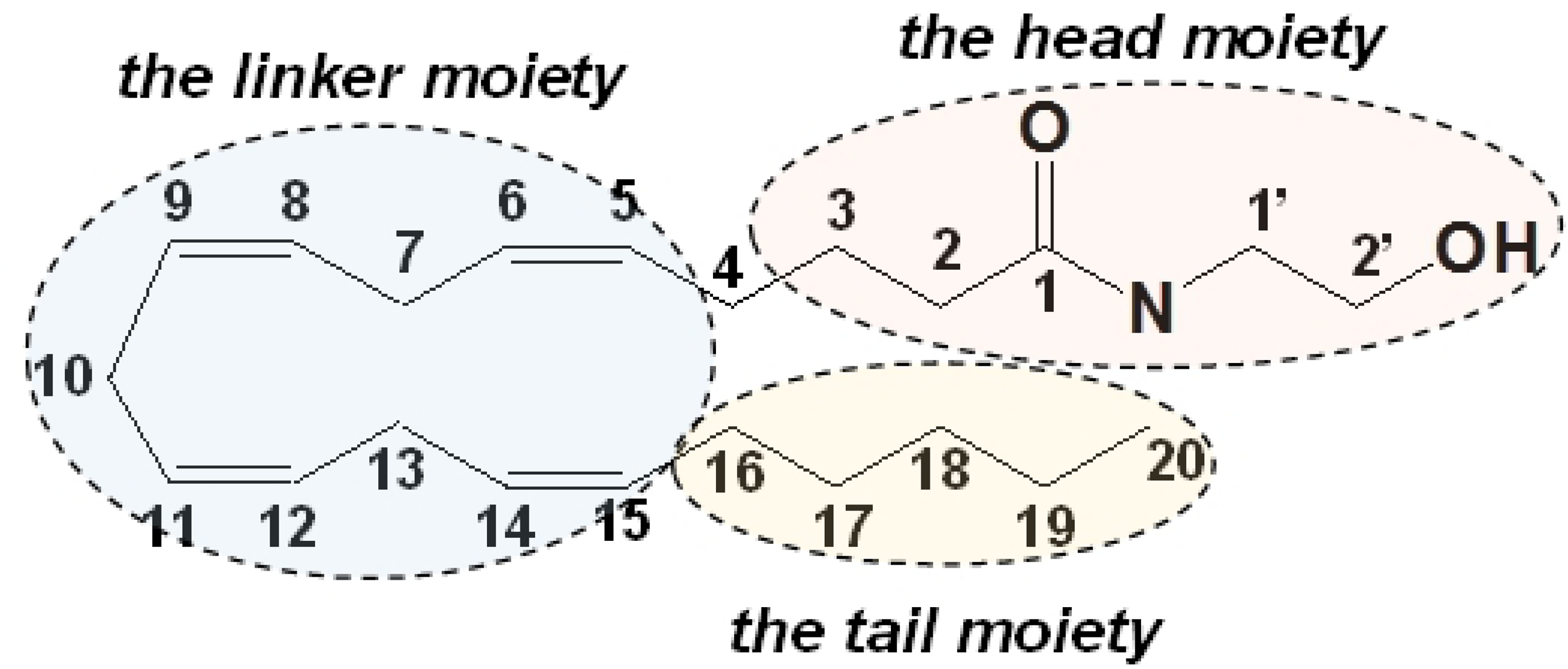
Structure of anandamide (AEA). The structure of AEA consists of three moieties, including the polar head moiety, the polyene linker moiety, and the hydrophobic tail moiety.

Our initial motivation of the present study was due to some intriguing results from recent studies demonstrating that the CB1 allosteric modulators (AMs) such as lipoxin A_4_ and ZCZ011 enhance selectively the AEA-activated CB1 receptors [18,19,20]. As the first step toward understanding how CB1 AMs allosterically enhance AEA-activated CB1 receptors, we felt imperative to determine the binding conformations of AEA responsible for CB1 receptor activation. In the present study, by using a combination of molecular docking and molecular simulations approaches, we explored many possible binding conformations of AEA within the binding pocket of the CB1 receptor and identified essential AEA binding domains. Our results indicate that the hydrophobic channel interactions are crucial for AEA binding to the CB1 receptor. Our results also suggest that the H2/H3 region of the CB1 receptor is an AEA binding subsite privileged possibly over the H7 region.

## Methods

### Determination of the AEA binding model

A low-energy ligand structure of AEA was obtained by performing the conformational analysis by using the MMFF molecular mechanics force field [21] implemented in the SPARTAN computational modeling package (Spartan’16, Wavefunction, Inc.). Initial docking poses of AEA were generated by using AutoDock4 [22]. For the receptor template, the CB1 receptor in the CB1-Gi complex model [23] refined according to the X-ray crystal structure of the AM11542-bound CB1 receptor [6] was used. The validity of the CB1 receptor model was partly confirmed by the overlay of the classical cannabinoid HU210 bound to the CB1 receptor in the refined CB1-Gi complex model to AM11542 in the X-ray crystal structure of the AM11542-bound CB1 receptor [6], which shows almost identical positions (see **Figure in** S1 Fig). For exploring AEA binding to the CB1 receptor, a grid box was created by setting 60 grid points in the x and y dimensions and 56 grid points in the z dimension with 0.375 Å spacing between grid points (i.e., a box of 22.5 Å x 22.5 Å x 21.0 Å) such that it covered the entire orthosteric binding pocket region. The position of the center of the grid box was guided by AM11542 bound to the CB1 receptor in the X-ray crystal structure of the AM11542-bound CB1 receptor [6]. A typical setting of docking parameters for performing AutoDock runs using a hybrid global-local Lamarkian genetic algorithm (LGA) [24] were: the rate of gene mutation (0.02), rate of crossover (0.8), GA window size (10), the number of individuals in population (150), the maximum number of energy evaluations in each run (25,000,000), the maximum number of generations (27,000) and the number of LGA docking runs (10). Only the ligand was allowed to freely move inside the grid box while the protein was rigidly fixed in position. The resulting docking poses were evaluated by the AutoDock4 scoring function [25]. AutoDock runs were performed more than one hundred times using the best scoring docking pose from the previous run as the starting pose for the next run. For every run the same grid box was used.

The best scoring docking poses obtained from the above AutoDock runs were overlaid to AM11542 bound to the CB1 receptor in the X-ray crystal structure [6]. Then, depending upon how AEA interacted with the hydrophobic channel, where the DMH chain of AM11542 was occupied in the X-ray crystal structure of the AM11542-bound CB1 receptor [6], they were clustered into three docking pose groups: 1) AEA docking pose Group **1** where the hydrophobic tail moiety of AEA occupied the hydrophobic channel; 2) AEA docking pose Group **2** where the polar head moiety of AEA occupied the hydrophobic channel; and 3) AEA docking pose Group **3** where the hydrophobic channel left unoccupied. Three representative poses were selected from AEA binding pose Group **1**. Similarly, three and two representative poses were selected from AEA docking pose Group **2** and AEA docking pose Group **3**, respectively. Overall, a total of eight docking poses were selected (Fig 2).

**Fig 2.**
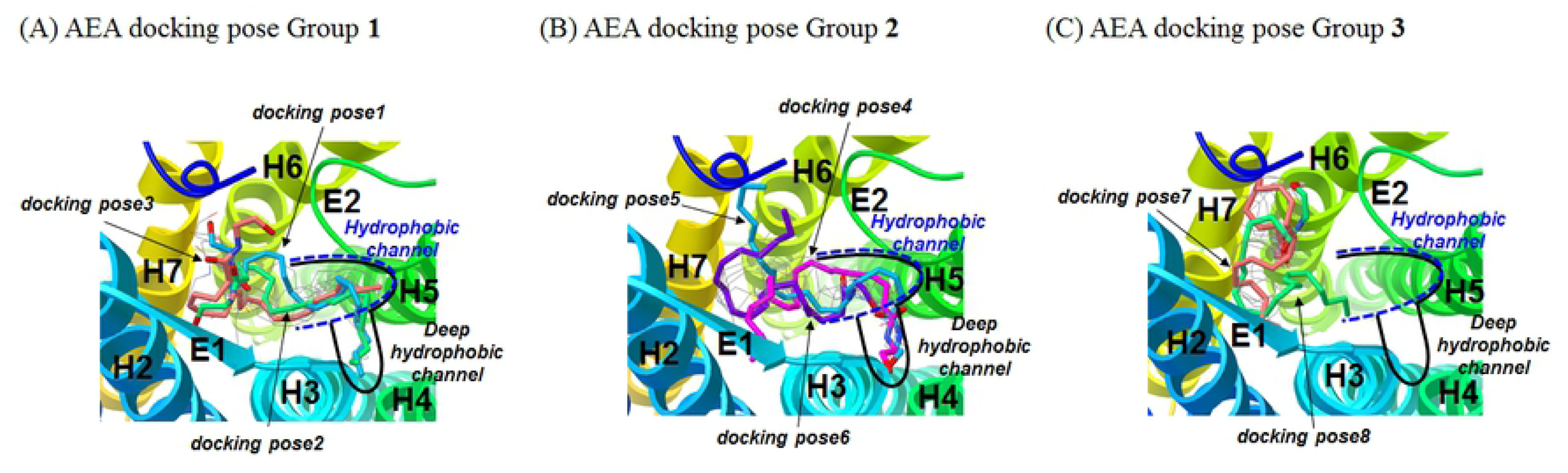
Selection of eight representative AEA docking poses from AutoDock docking poses. (A) AEA docking pose Group **1**: *docking pose1* (in cyan), *docking pose2* (in green) and *docking pose3* (in orange). (B) AEA docking pose Group **2**: *docking pose4* (in magenta), *docking pose5* (in cyan) and *docking pose6* (in purple). (C) AEA docking pose Group: *docking pose7* (in orange) and *docking pose8* (in green). Not all of the Autodock docking poses are displayed for clarity.

### Molecular dynamics simulations of the CB1-G assembly

Each of the eight selected AEA docking poses inserted into the binding pocket of the CB1 receptor in the CB1-Gi complex model in a fully hydrated 1-palmitoyl-2-oleoyl-*sn*-glycero-3-phosphocholine (POPC) lipid bilayer was subjected to energy minimization (5,000 iterations). This was followed by a molecular dynamics (MD) simulation at 310 K in the NPT ensemble to obtain an all-atom, solvent-equilibrated AEA binding pose. A long time (typically 200 ns) MD simulation of each of the eight selected docking poses was performed to ascertain receptor binding interaction of AEA became optimally established as indicated by the RMSDs of the receptor as well as the bound ligand AEA.

### Simulation Protocol

All simulations were performed using the NAMD simulation package (ver. 2.7 Linux-x86_64) [26], using CHARMM36 force field parameters for proteins with the ϕ/Ψ angle cross-term map correction [27,28] and lipids [29], and the TIP3P water model [30]. The temperature was maintained at 310 K through the use of Langevin dynamics [31] with a damping coefficient of 1/ps. The pressure was maintained at 1 atm by using the Nosé-Hoover method [32] with the modifications as described in the NAMD user’s guide. The van der Waals interactions were switched at 10 Å and zero smoothly at 12 Å. Electrostatic interactions were treated using the Particle Mesh Ewald (PME) method [33]. A pair list for calculating the van der Waals and electrostatic interactions was set to 13.5 Å and updated every 10 steps. A multiple time-stepping integration scheme, the impulse-based Verlet-I reversible reference system propagation algorithm method [34], was used to efficiently compute full electrostatics. The time step size for integration of each step of the simulation was 1 fs.

### CHARMM parameterization

To describe AEA in the MD simulations using the CHARMM force field, missing parameters were determined. To minimize any inconsistency with the existing CHARMM parameters, most of the missing parameters for describing AEA were borrowed from the parameter values of lipids and chemically relevant structures.

### RMSD analysis

RMSD values of the CB1 receptor were calculated by root mean square fitting to the initial coordinates with respect to the backbone Cα atoms of the TM helical residues of the CB1 receptor (TM1: Pro113-His143; TM2: Tyr153-His178; TM3:Arg186-Ser217; TM4:Arg230-Val249; TM5: Glu273-Val306; TM6: Met337-Ile362; TM7:Lys373-Arg400). The RMSD values of the polar head moiety of AEA bound to the above fitted CB1 receptor were calculated with respect to the initial coordinates after fitting to the heavy atoms of its head moiety (Fig 1). Similarly, the RMSD values of the hydrophobic tail moiety of AEA bound to the above fitted CB1 receptor are calculated with respect to the initial coordinates after fitting to the heavy atoms of the hydrophobic tail moiety (Fig 1).

## Results

### Eight representative AEA docking poses selected from AutoDock docking runs

The eight representative docking poses selected from more than one hundred AutoDock docking runs are shown in Fig 2. These docking poses were clustered into AEA docking pose Group **1**, AEA docking pose Group **2**, and AEA docking pose Group **3**, depending upon how AEA interacted with the hydrophobic channel or the deep hydrophobic channel (In our AutoDock docking runs, AEA sometimes occupied the hydrophobic channel deeper than AM11542 in the X-ray crystal structure of AM11542-bound CB1 receptor [6]. Thus, this extended hydrophobic channel was called the deep hydrophobic channel.)

In three selected poses (named *docking pose1, docking pose2* and *docking pose3*) that belong to AEA docking pose Group **1**, the tail moiety of AEA commonly occupied the hydrophobic channel or the deep hydrophobic channel (Fig 2A). In *docking pose1*, the tail moiety of AEA occupied the deep hydrophobic channel and the head moiety of AEA bound the H7 region. Thus, *docking pose1* was assigned to be docking pose Group **1d_H7** (“**1d**” denotes that the tail moiety binds the ***d****eep* hydrophobic channel and “**H7**” denotes the H7 region where the head moiety binds). In *docking pose2*, the tail moiety occupied the ***d****eep* hydrophobic channel and the head moiety bound the H2/H3 region. Thus, *docking pose2* was assigned to be docking pose Group **1d_H2/H3**. In *docking pose3*, the tail moiety occupied the hydrophobic channel and the head moiety bound the H7 region. Thus, *docking pose3* was assigned to be docking pose Group **1_H7**.

For the three selected docking poses (named *docking pose4, docking pose5* and *docking pose6*) that belong to AEA docking pose Group **2**, the head moiety of AEA commonly occupied the hydrophobic channel or the deep hydrophobic channel (Table 1). In *docking pose4*, the head moiety of AEA occupied the deep hydrophobic channel and the tail moiety bound the H2/H3 region (Fig 2B). Thus, *docking pose4* was assigned to be **2d_H2/H3** (“**2d**” denotes that the head moiety binds the ***d****eep* hydrophobic channel and “**H2/H3**” denotes the H2/H3 region where the tail moiety binds). In *docking pose5*, the head moiety occupied the ***d****eep* hydrophobic channel and the tail moiety bound the H7 region. Thus, *docking pose5* was assigned to be **2d_H7**. In *docking pose6*, the head moiety occupies the hydrophobic channel and the tail moiety points toward the middle of the binding core toward the EC region (i.e., the pocket outer core). Thus, *docking pose6* was assigned to be **2_OC** (“**2**” denotes Group **2** and “**OC**” denotes the outer core region).

**Table 1.** Receptor interactions of the eight docking poses selected from AutoDock runs and the corresponding equilibrated poses in simulation.

| AEA pose | The orthosteric binding pocket |  |  |  | Group |
| --- | --- | --- | --- | --- | --- |
|  | Hydrophobic channel | Deep hydrophobic channel | H2/H3 region | H7 region |  |
| *Docking pose1* |  | tail |  | head | **1d_H7** |
| *Equilibrated pose1* | tail |  |  | head | **1_H7** |
| *Docking pose2* |  | tail | head |  | **1d_H2/H3** |
| *Equilibrated pose2* | tail |  | head |  | **1_H2/H3** |
| *Docking pose3* | tail |  |  | head | **1_H7** |
| *Equilibrated pose3* | tail |  |  | head | **1_H7** |
| *Docking pose4* |  | head | tail |  | **2d_H2/H3** |
| *Equilibrated pose4* |  | head | tail |  | **2d_H2/H3** |
| *Docking pose5* |  | head |  | tail | **2d_H7** |
| *Equilibrated pose5* |  | head | tail |  | **2d_H2/H3** |
| *Docking pose6*<sup>a)</sup> | head |  |  |  | **2_OC** |
| *Equilibrated pose6* |  | head | tail |  | **2d_H2/H3** |
| *Docking pose7*<sup>a),b)</sup> |  |  |  |  | **3_IC_H2/H3** |
| *Equilibrated pose7* |  | head | tail |  | **2d_H2/H3** |
| *Docking pose8*<sup>b)</sup> |  |  |  | head | **3_H7_IC** |
| *Equilibrated pose8* | tail |  |  | head | **1_H7** |

<sup>a)</sup>The tail moiety points toward the middle of the binding core toward the EC region (i.e., the pocket outer core).
<sup>b)</sup>The head moiety points toward deep inside the binding core toward the IC region (i.e., the pocket inner core).

For the two selected docking poses (named *docking pose7* and *docking pose8*) that belong to AEA binding pose Group **3**, the hydrophobic channel or the hydrophobic channel was commonly left unoccupied (Fig 2C). In *docking pose7*, the tail moiety bound the H2/H3 region and the head moiety points toward deep inside the binding core toward the IC region (i.e., the pocket inner core). Thus, *docking pose7* was assigned to be **3_IC_H2/H3** (“**3**” denotes Group 3, “**IC**” denotes the outer core region where the head moiety binds, and “**H2/H3**” denotes the H2/H3 region where the tail moiety binds). In *docking pose8*, the head moiety bound the H7 region and the tail moiety binds the pocket inner core. Thus, *docking pose8* was assigned to be **3_H7_IC** (“**3**” denotes Group 3, “**H7**” denotes the H7 region where the head moiety binds, and “**IC**” denotes the inner core region where the tail moiety binds).

### RMSD analysis of the eight representative AEA docking poses

As shown in **Figure in** S2 Fig, all the receptors in the AEA-bound CB1-G complex model systems were converged with the r.m.s.d. values <2 Å with respect to the Cα atoms of the receptor TM helical bundle, indicating the CB1 receptor models became stable at the end of the simulation. Similarly, all the ligands in the AEA-bound CB1-G complex model systems were converged at the end of the simulation, indicating the bound ligand became stable. The RMSD values of the head moiety and the tail moiety of the bound ligand in the eight AEA docking poses are shown in Fig 3. The RMSD values of the tail moiety of AEA in *docking pose3* and *docking pose7* and the RMSD values of the polar head moiety of AEA in *docking pose3, docking pose4, docking pose5*, and *docking pose8* showed little change, indicating that these binding interactions remained stable throughout the simulation. However, significant increases in the RMSD values of the head moiety or the tail moiety indicated that the regions initially occupied by the head moiety or the tail moiety became noticeably altered in simulation. Significant increases in the RMSD values are seen for the hydrophobic tail moiety of AEA in *docking pose1* and *docking pose2* (a shift out from the deep hydrophobic channel to the hydrophobic channel), *docking pose5* and *docking pose6* (a shift from the H7 region to the H2/H3 region) and *docking pose8* (a shift from the binding pocket core region to the hydrophobic channel) (Fig 3). Similarly, significant increases in the RMSD values are seen for the polar head moiety of AEA in *docking pose2* (a shift from the binding pocket core region to the H2/H3 region), *docking pose6* (a shift from to the hydrophobic channel to the deep hydrophobic channel), *docking pose7* (a shift from the binding pocket core region to the deep hydrophobic channel) (Fig 3). The eight docking poses and equilibrated poses are summarized in Table 1.

**Fig 3.**
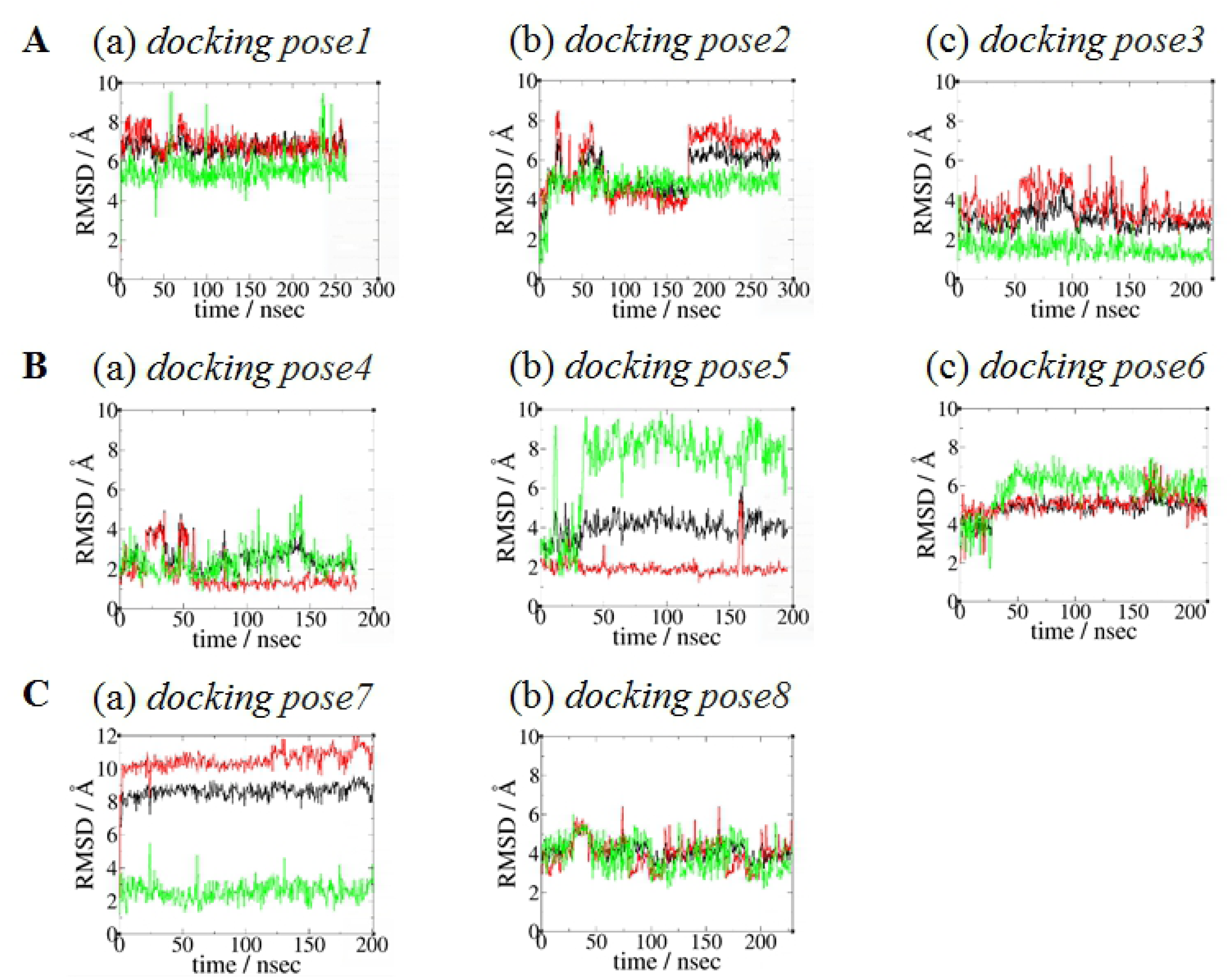
RMSD plots of the eight AEA docking poses. (A) RMSD plots of AEA docking pose Group **1**. (B) RMSD plots of AEA docking pose Group **2**. (C) RMSD plots of AEA docking pose Group **3**. The r.m.s.d. values of the whole molecule (in black), the polar head moiety (in red), and the hydrophobic tail moiety (in green) of the bound AEA (in red) in eight AEA docking poses, calculated with respect to the initial coordinates (heavy atoms only) after fitting the proteins based upon the backbone heavy atoms of the TM helical residues of the CB1 receptor.

### Three AEA binding poses merged from eight equilibrated poses

As shown in Fig 4, the equilibrated poses share the binding region in the orthosteric binding pocket well with the cannabinoid ligands in the orthosteric binding pocket as found in the X-ray crystal structures of the AM11542-bound CB1 receptor [6] and the CP55940-bound CB1 receptor [7]. Each of the eight equilibrated poses adopts an extended conformation that spans throughout the binding core region. It is also revealed in all of the equilibrated poses that the hydrophobic channel is always occupied by the tail moiety or the head moiety of AEA. As shown in Fig 5, most of the equilibrated poses are quite different from their initial docking poses in conformation and position (Table 1). This can be also seen in the RMSD plots of these docking poses (Fig 3).

**Fig 4.**
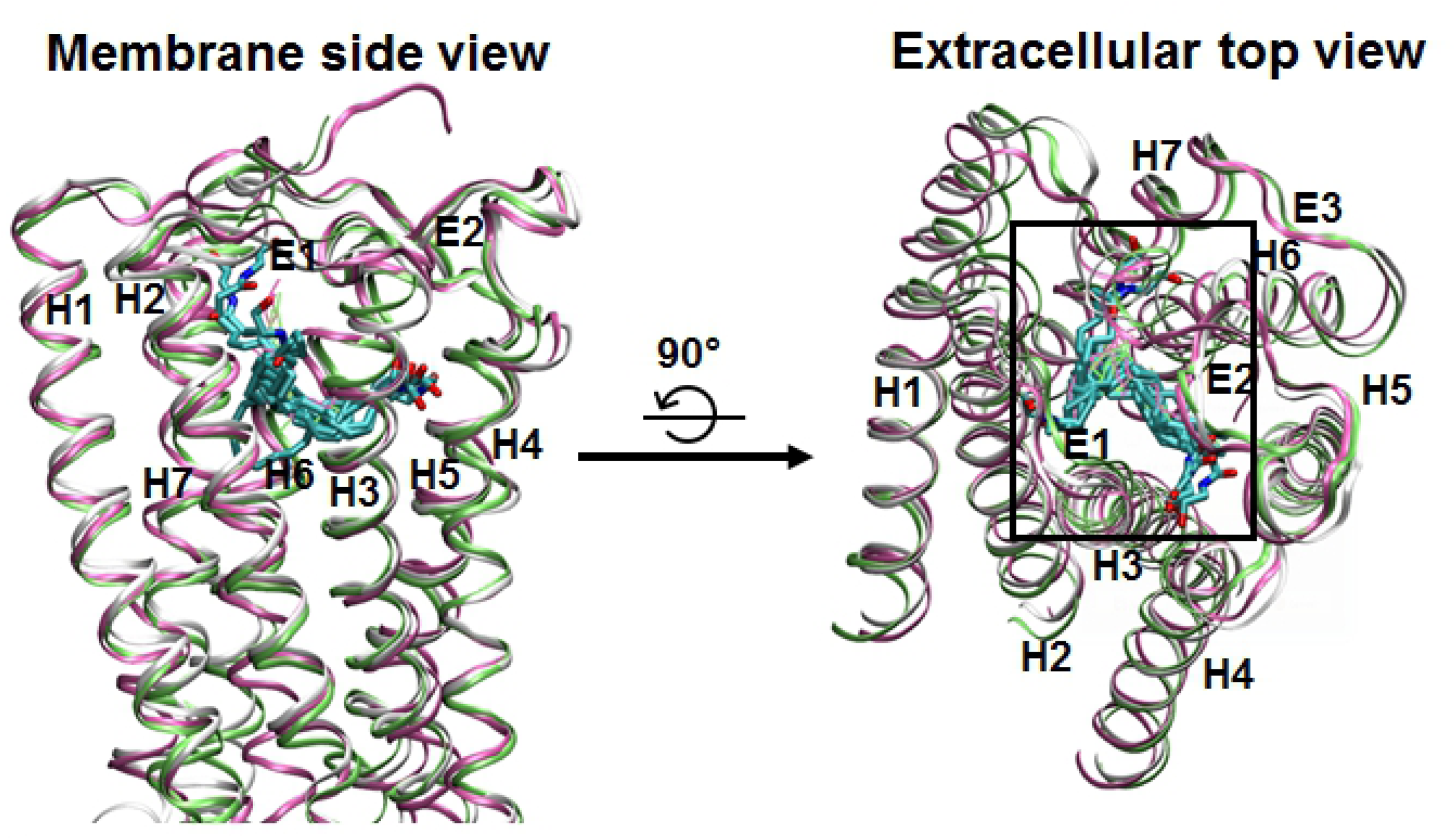
Superposition of the CB1 receptor in AEA *equilibrated pose1* and the X-ray crystal structures of the CB1 receptor. All the eight equilibrated poses (in atom type) are overlaid to AM11542 (in green) and CP55940 (in mauve) after the receptor in these poses were superimposed to the receptors in the X-ray crystal structures of the AM11542-bound CB1 receptor [6] and the CP55940-bound CB1 receptor [7]. The alignment rule: the backbone Cα atoms of H3 (Arg186-Ser217), H4 (Arg230-Val249), H5 (Glu273-Val306) and H7 (Lys373-Arg400). Color coding: carbon, cyan; oxygen, red; and nitrogen, blue. Hydrogen atoms are omitted for clarity.

**Fig 5.**
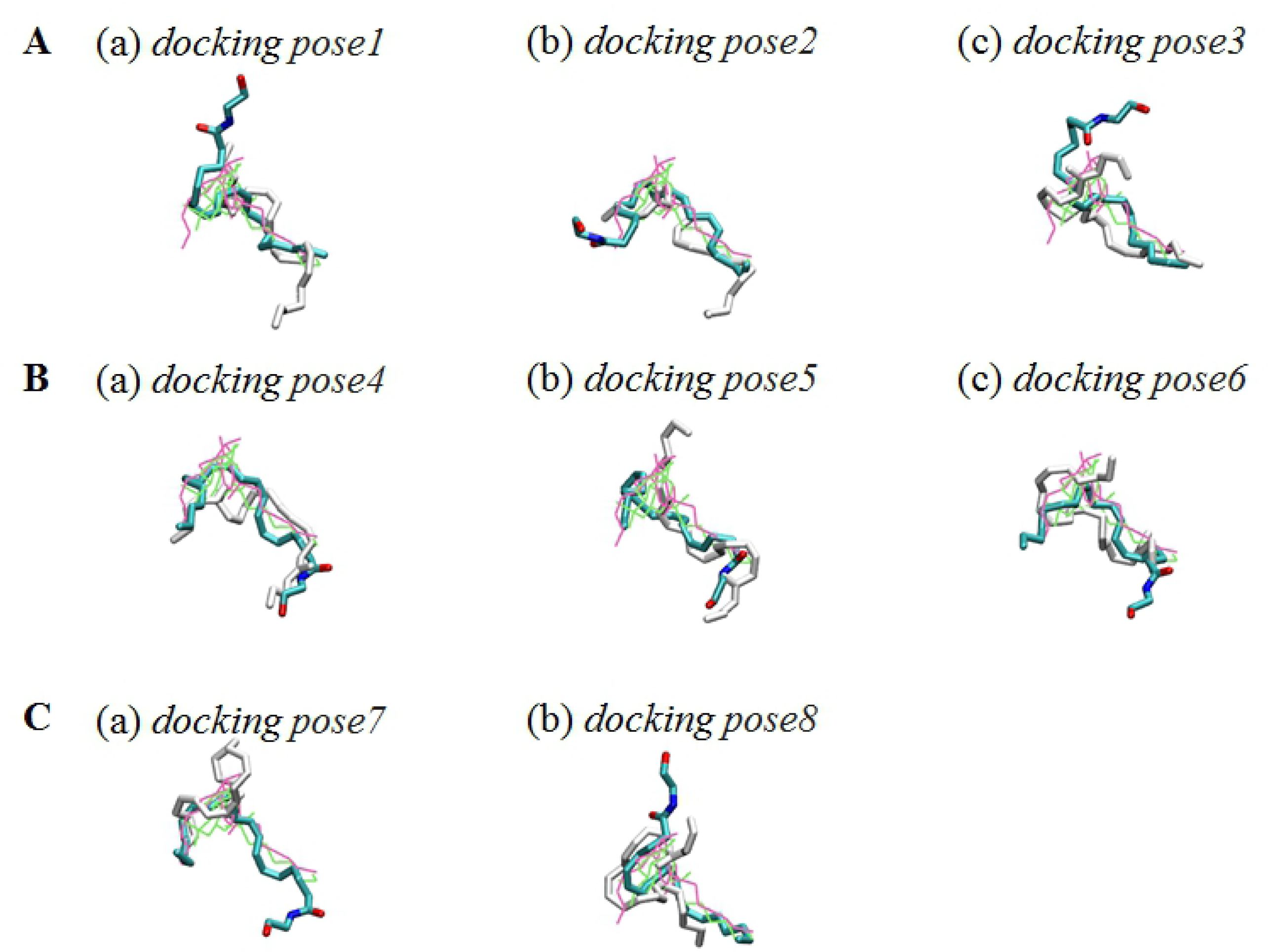
Overlay of the eight docking poses and equilibrated poses. (A) AEA docking pose Group **1** (*docking pose1, docking pose2*, and *docking pose3*). (B) AEA docking pose Group **2** (*docking pose4, docking pose5*, and *docking pose6*). (C) AEA docking pose Group **3** (*docking pose7* and *docking pose8*). The eight docking poses (in white) and equilibrated poses (in atom type) were superimposed to AM11542 (in green) and CP55940 (in mauve) in the X-ray crystal structures of AM11542-bound CB1 receptor [6] and CP55940-bound CB1 receptor [7]. The alignment rule of the receptors is same as in Fig 4. Color coding: carbon, cyan; oxygen, red; and nitrogen, blue. Hydrogen atoms are omitted for clarity.

After overlaid to the cannabinoid ligands in the orthosteric binding pocket as found in the X-ray crystal structures of the AM11542-bound CB1 receptor [6] and the CP55940-bound CB1 receptor [7], the eight equilibrated poses became merged into **1_H7, 1_H2/H3** and **2d_H2/H3** (Fig 6). No AEA equilibrated pose is found such that the tail moiety of AEA binds the deep hydrophobic channel (Figs 6A and 6B). Also, no AEA equilibrated pose is found such that the tail moiety binds the H7 region when the head moiety binds to the deep hydrophobic channel (i.e., **2_H7**).

**Fig 6.**
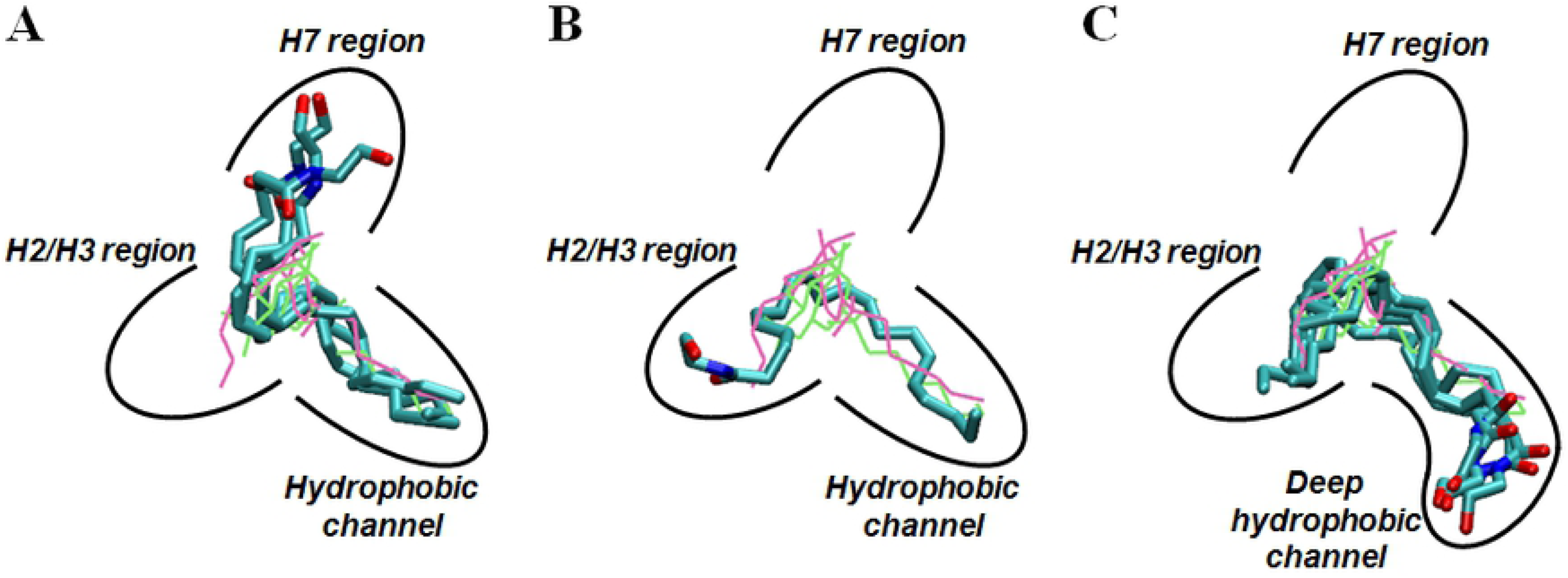
Three AEA equilibrated poses. (A) AEA equilibrated pose **1_H7** (*equilibrated pose1, equilibrated pose3* and *equilibrated pose8*). (B) AEA equilibrated pose **1_H2/H3** (*equilibrated pose2*). (C) AEA equilibrated pose **2d_H2/H3** (*equilibrated pose4, equilibrated pose5, equilibrated pose6* and *equilibrated pose7*).

Detailed structural features of each of three AEA equilibrated poses are described below: a) Equilibrated pose **1_H7**. *Equilibrated pose1, equilibrated pose3* and *equilibrated pose8* are merged into the equilibrated **1_H7** (Table 1 and Fig 6A). The hydrophobic tail moiety is well aligned with the DMH chain of AM11542 and CP55940 and the head moiety binds the H7 region. It is interesting to see that not only the terminal five carbons (C16-C20) but also the fourth double bond (C14=C15) binds the hydrophobic channel (Figs 6A and 6B). The detailed receptor interactions of AEA in *equilibrated pose1* are shown in Fig 7A. The tail moiety of AEA occupies the hydrophobic pocket formed by Leu193, V196, Thr197, Phe200, Ile271, Phe268, Tyr275, Leu276, Trp279, Leu359, and Met363. The head moiety of AEA is surrounded by Cys107, Phe108, His178, Lys376, Phe379, and Ala380. The middle linker moiety in equilibrated pose **1_H7** interacts with a group of aromatic residues in the binding core, including Phe103, Phe174, Phe177, His178, Phe189, Phe268, and Phe379, through aromatic-π stacking interactions (see **Figure in** S3A Fig).

**Fig 7.**
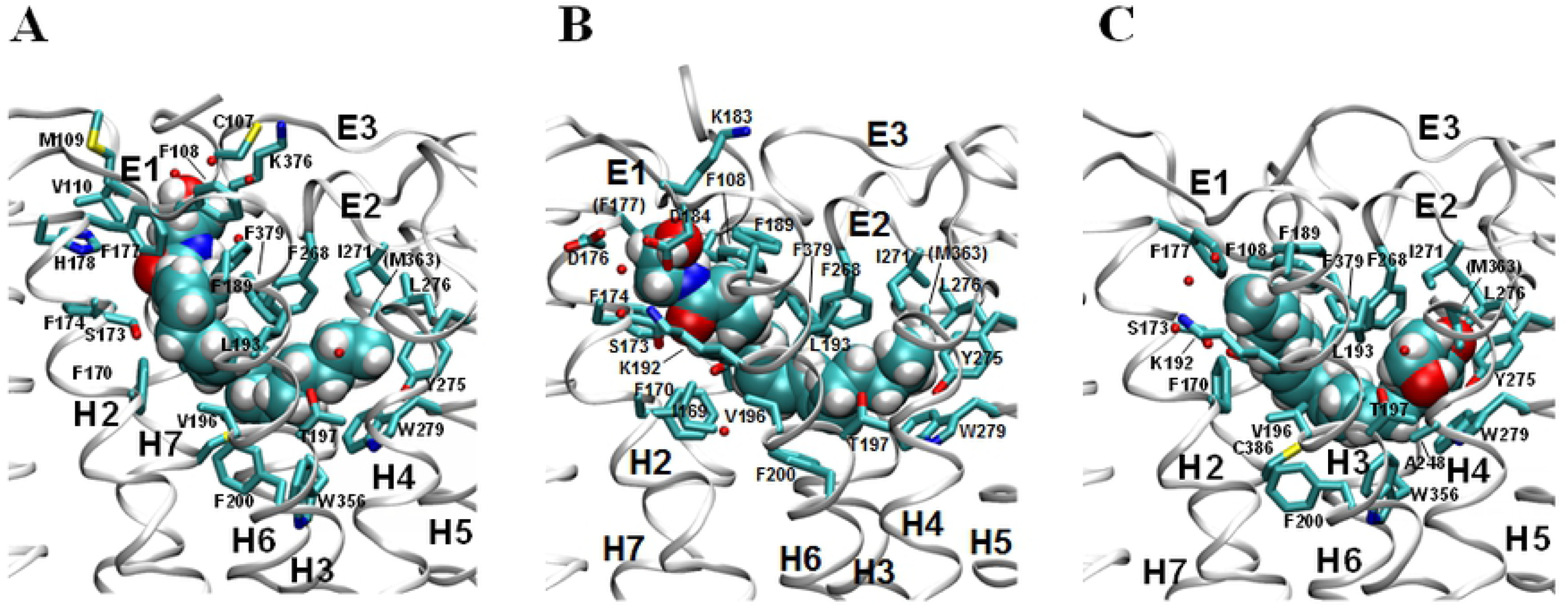
Detailed receptor binding interactions of three AEA equilibrated poses. (A) AEA equilibrated pose **1_H7** (*equilibrated pose1*). (B) AEA equilibrated pose **1_H2/H3** (*equilibrated pose2*). (C) AEA equilibrated pose **2d_H2/H3** (*equilibrated pose4*). b) Equilibrated pose **1_H2/H3**. *Equilibrated pose2* remains as the equilibrated pose **1_H2/H3** (Table 1 and Fig 6B). The tail moiety occupies the hydrophobic channel just as in the equilibrated pose **1_H7**, while the head moiety binds the H2/H3 region. The detailed receptor interactions of AEA in *equilibrated pose2* are shown in Fig 7B. The head moiety closely interacts with D184 and Lys192. The hydroxyl group of the head moiety is H-bond to Asp184, which forms a salt bridge with Lys192 (see **Figure in** S4 Fig). In addition, Asp184 forms water-mediated H bonds to Asp176. It appears that the extensive H-bonding network centered at Asp184 and Lys192 contributes favorably to the binding of the polar head moiety of AEA. The head moiety binds the H2/H3 region under E1 extensively, including Ile169, Ser173, Phe174, Asp176, Phe177, Lys183, Phe189, Lys192, and Leu193. The middle linker moiety in the equilibrated pose **1_H2/H3** interacts with a group of aromatic residues in the binding core through aromatic-π stacking interactions similar to the equilibrated pose **1_H7** (see **Figure in** S3B Fig). c) Equilibrated pose **2d_H2/H3**. *Equilibrated pose4, equilibrated pose5, equilibrated pose6* and *equilibrated pose7* are merged into the equilibrated pose **2d_H2/H3** (Table 1 and Fig 6C). In this equilibrated pose, the polar head moiety binds to the deep hydrophobic channel while the tail moiety binds to the H2/H3 region. The detailed receptor interactions of AEA in *equilibrated pose4* are shown in Fig 7C. Most of the binding residues that interact with the tail moiety are somewhat similar to not as extended as those that interact with the tail moiety in **1_H2/H3**. The binding residues that interact with the head moiety are similar to those in the equilibrated pose **1_H7** or **1_H2/H3**. However, the terminal hydroxyethyl group of the head moiety occupies the deep hydrophobic pocket (Fig 7C). A close examination reveals that the N atom of the amide moiety forms an H-bond to Thr197, while the O atom of the hydroxyethyl group frequently forms an H-bond to Tyr275. It appears that these H-bonds to the polar residues play key roles in stabilizing the polar head moiety deep inside the hydrophobic channel. Similar to the equilibrated poses **1_H7** and **1_H2/H3**, the linker moiety in the equilibrated pose **2d_H2/H3** interacts with a group of aromatic residues in the binding core through aromatic-π stacking interactions (see **Figure in** S3C Fig).

## Discussion

### The importance of the hydrophobic channel in AEA binding to the CB1 receptor

All of the eight representative equilibrated AEA binding poses show that AEA interacts tightly with the hydrophobic channel. If either the tail moiety or the head moiety initially binds to the hydrophobic channel, it remained there throughout the simulation (Figs 3A, 3B, 5A and 5B). If the hydrophobic channel initially left empty, it became occupied by either the tail or the head moiety in simulation (Figs 3C and 5C). These results underscore the importance of the hydrophobic channel of the CB1 receptor in AEA binding just as seen in the recent X-ray crystal structures of the CB1 receptor in complex with various ligands [4,6,7,35]. In support, it has been reported that Ala mutations of the hydrophobic channel forming residues Leu193 and Met363 of the CB1 receptor caused ∼80-fold and ∼4-fold decreases in CP55940 binding [36]. Since the tail moiety of AEA in the equilibrated poses **1_H7** and **1_H2/H3** is exactly overlaid to the DMH alkyl chain of AM11542 [6] and CP55940 [7], mutation of these hydrophobic pocket residues would also alter AEA binding affinity.

It is surprising to see in the present study that the polar head moiety of AEA is also able to stably occupy the hydrophobic channel (as in the equilibrated pose **2d_H2/H3**) (Fig 6C). It appears that the stabilization of the polar head moiety through H-bonding is required for its binding to the deep hydrophobic channel.

### Which binding region is a privileged subsite?

It is shown from the present study that regardless of whether the hydrophobic pocket was occupied or empty in the initial docking poses, the hydrophobic channel become preferentially occupied in all of the eight equilibrated poses in simulation. Therefore, it is likely that if the one end moiety (either the head moiety or the tail moiety) of AEA establishes its binding interaction with the hydrophobic channel as the primary binding contact, then the other end moiety of AEA establishes its binding to either the H2/H3 region or the H7 region before the linker moiety forced to be conformationally much restricted to complete AEA binding to the receptor.

The recently determined X-ray crystal structure of the classical cannabinoid agonist AM11542-bound CB1 receptor [6] reveals that the trimethyl substituted B/C-ring moiety of AM11542 binds the H2/H3 region. Similarly, the X-ray crystal structure of the nonclassical CP55940-bound CB1 receptor [7] reveals that the propylhydroxyl substituted C-ring moiety of CP55940 bind preferentially the H2/H3 region. A 10-fold increase in binding affinity by the introduction of the propylhydroxyl group to the C-ring of CP47497, which becomes equivalent to CP55940 [37], supports the idea that the H2/H3 region is important for cannabinoid binding. Alanine mutations of the H2/H3 residues Phe174, Phe177, Asp184, Phe189, Lys192 and Leu193 resulted in significant decreases in binding affinity of CP55940 [8,36,38,39], also underscoring the importance of the H2/H3 region in cannabinoid binding.

The equilibrated pose **1_H2/H3** uniquely shows extensive H2/H3/E1 interactions of the head moiety of the ligand (see **Figure in** S4 Fig). Collectively, these experimental results indicate that the H2/H3 region of the CB1 receptor is a ligand binding subsite privileged over the H7 region. In the present study, AEA interacts with the H2/H3 region in the equilibrated poses **1_H2/H3** and **2d_H2/H3**, while AEA little interacts with the H2/H3 region in the equilibrated pose **1_H7**. As shown in **Table in** S1 Table, the CB1-AEA binding interaction was estimated by the nonbonding interaction energy between the binding pocket residues and the bound ligand AEA. Based on the estimated binding interaction energy values, it is predicted that the equilibrated pose **1_H2/H3** interact with the receptor more strongly than the equilibrated pose **1_H7**. Because the tail moiety of the ligand in both the equilibrated pose **1_H2/H3** and **1_H7** interacts with the hydrophobic channel quite similarly (see Figs 6 and 7), the favorable binding interaction shown in the equilibrated pose **1_H2/H3** over the equilibrated pose **1_H7** suggests that AEA interactions with the H2/H3 region is more important than with the H7 region.

### Which AEA binding pose is the best candidate for the bioactive conformation?

If we assume that all of the equilibrated poses **1_H7, 1_H2/H3** and **2d_H2/H3** as potential candidates for the bioactive conformation, the measured binding affinity of AEA to the CB1 receptor would be the results of the binding of these poses in equilibrium. If the equilibrated poses **1_H7** and **2d_H2/H3** are weaker binding modes than the equilibrated pose **1_H2/H3**, as predicted by the estimated the CB1-AEA binding interaction (**Table in** S1 Table), some ligand binding exerted by the equilibrated poses **1_H7** and **2d_H2/H3** may still be present but would be weaker than ligand binding exerted by the equilibrated pose **1_H2/H3**. In this regard, an increase in CB1 affinity by substituting the 2-hydroxyethyl group of AEA with a cyclopropyl ring or a halogen [41-43] is intriguing. It is possible that such a hydrophobic substitution for the polar head moiety of AEA would make the equilibrated pose **2d_H2/H3** a more favorable binding mode, contributing to an increase in AEA binding affinity overall.

Compared with the equilibrated pose **2d_H2/H3**, the equilibrated pose **1_H2/H3** exhibits extensive binding interactions with the H2/H3 region under E1 (Fig 7B), including an H-bond to Asp184 (**Figure in** S4 Fig). On the other hand, the binding interactions with the hydrophobic channel in the equilibrated pose **1_H2/H3** presumably less extensive than the binding interactions with the deep hydrophobic channel in the equilibrated pose **2d_H2/H3** (Fig 7C). Therefore, it is expected that the overall binding interactions in the equilibrated poses **1_H2/H3** and **2d_H2/H3** would be quite competitive. However, the estimated the CB1-AEA binding interaction energy values predict that the equilibrated pose **1_H2/H3** binds the receptor much stronger (∼ 10 kcal/mol) than the equilibrated pose **2d_H2/H3** (**Table in** S1 Table), suggesting that in the equilibrated pose **2d_H2/H3** the binding interactions of the polar head moiety with the deep hydrophobic channel is not advantageous for compensating for the limited binding interactions with the H2/H3 region of the tail moiety. In agreement, our simulation results of *docking pose1* and *docking pose2* reveal that the binding interactions of the tail moiety of AEA with the deep hydrophobic channel is not favored (Fig 3A). Moreover, the equilibrated pose **2d_H2/H3** would not be a plausible binding mode in physiological environments, because the polar head moiety of AEA would not easily reach the hydrophobic channel located deep inside the binding pocket. Overall, it is less likely that the equilibrated pose **2d_H2/H3** is an ideal candidate for the bioactive conformation of AEA.

On the other hand, the equilibrated pose **1_H2/H3** could be a better candidate for the bioactive conformation than the equilibrated pose **1_H7**, in consideration of the recent X-ray crystal structures of the CB1 receptor [6,7] and the available mutational studies suggesting the H2/H3 region of the CB1 receptor offers a binding subsite privileged over the H7 region. Overall, the equilibrated pose **1_H2/H3** could be the best candidate for the bioactive conformation of AEA. In order to check the validity of the equilibrated pose **1_H2/H3**, another independent MD simulation was carried out, starting from a docking pose (named *docking pose2’*) different from *docking pose2* within AEA docking pose group **1_H2/H3**. The resulting *equilibrated pose2’* was quite similar to *equilibrated pose2* (**Figure in** S5 Fig).

The chance of the bioactive conformation being present in AEA is much lower than in AM11542 and CP55940, simply because AEA is structurally far more flexible than AM11542 and CP55940. It is difficult for the highly flexible AEA to be locked into the active conformation required for best fitting to the binding pocket. Both the varying polar head moiety and the varying hydrophobic tail of AEA would interfere significantly from achieving the bioactive conformation. Overall, AEA is expected to achieve the active conformation much more difficult than AM11542 and CP55940, possibly contributing to its known partial agonistic activity [1].

## Conclusions

In summary, we have explored many possible binding conformations of AEA within the binding pocket of the CB1 receptor well defined in the recently determined X-ray crystal structures of the ligand-bound CB1 receptors, by using a combination of docking and MD simulation approaches. Because the challenging problem of the conformational explosion in AEA was significantly reduced owing to the binding preference of AEA to the hydrophobic channel, we were able to predict essential AEA binding domains of the CB1 receptor. Although the present study is rather limited in exploring all the available binding conformations allowed for the extremely flexible AEA, our results suggest that CB1 receptor interactions of the H2/H3 region as well as the hydrophobic channel are important for AEA binding.

## Acknowledgments

I gratefully acknowledge Dr. Randall Griffus for his generous support for securing Hewlett Packard Z440 workstations with NVIDIA Tesla K40 GPUs and a Hewlett Packard Z4 G4 workstation with NVIDIA GP100 GPUs. I thank Mr. Alvaro Cortez, Ms. Arianna Fisher, and Ms. Nashely Hernandez for their critical reading and comments on the manuscript.

## Supporting information

**S1 Fig. Overlay of HU210 and AM11542**. HU210 (in gray) in the binding pocket of the CB1-Gi complex model [1], refined according to the X-ray crystal structure of the AM11542-bound CB1 receptor [2], is overlaid to AM11542 (in green) in the X-ray crystal structure of the AM11542-bound CB1 receptor [2]. The binding pocket residues within 4 Å of the ligand are also displayed.

**S2 Fig. The r.m.s.d. values of the CB1 receptor in the eight AEA docking poses.** The r.m.s.d. values were calculated by root mean square fitting to the initial coordinates with respect to the backbone heavy atoms of the TM helical residues of the CB1 receptor. (A) AEA docking pose Group **1**. (B) AEA docking pose Group **2**. (C) AEA docking pose Group **3**.

**S3 Fig. Aromatic-π Stacking interactions of the polyene linker moiety of AEA.** (A) Equilibrated pose **1_H7**. (B) Equilibrated pose **1_H2/H3**. (C) Equilibrated pose **2d_H2/H3**.

**S4 Fig. Key receptor interactions of the polar head moiety of AEA in the equilibrated pose 1_H2/H3.** Hydrogen bonding interactions are shown in red dotted lines. Hydrogen bonding distance (in Å) is also shown. Residues and water molecules are shown in stick mode and AEA are shown in space-filling mode.

**S5 Fig. Analysis of docking pose2’.** (A) *Docking pose2’* (in dark green) and the AEA docking poses (in line mode) that belong to the same cluster as *docking pose2* (in pink). (B) AEA equilibrated pose **1_H2/H3** (*equilibrated pose2* and *equilibrated pose2’*). (C) The r.m.s.d. values of the CB1 receptor in *docking pose2’*. (D) The RMSD plots of the head moiety and the tail moiety of AEA in *docking pose2’*. The r.m.s.d. values of the polar head moiety (in red) and the hydrophobic tail moiety (in green) of the bound AEA (in red), calculated with respect to the initial coordinates (heavy atoms only) after fitting the proteins based upon the backbone heavy atoms of the TM helical residues of the CB1 receptor.

**S1 Table. The CB1-AEA binding interaction estimated by the nonbonding interaction energy.** NAMD Energy Plugin as implemented in VMD [4] was used to calculate the nonbonding interaction energy values between the bound ligand AEA and any binding pocket residue. For the overall binding interaction energies, the whole ligand atoms were considered. For the head moiety of AEA, the ethanolamide and the propyl (C2-C4) atoms were used. For the polyene moiety of AEA, only the polyene (C5-C15) atoms were used. For the tail moiety of AEA, the terminal pentyl (C16-C20) atoms were used. A smooth switching function was activated at the distance of 10 Å to truncate the nonbonding interaction energies smoothly at the cutoff distance of 12 Å. The energy values with the standard deviation of the values in parentheses were averaged over the last 25.0 ns of the simulation.

